# Novel linkage disequilibrium-based genotype-by-environmental interaction method for genomic prediction of cotton yield and fibre quality traits

**DOI:** 10.64898/2026.05.03.722538

**Authors:** Zitong Li, Xuesong Li, Shiming Liu, Iain Wilson, Qian-Hao Zhu, Warwick Stiller, Warren Conaty

## Abstract

Genomic prediction (GP) across diverse environments has a potential to accelerate genetic gain in cotton breeding programs. A major challenge in GP is modelling genotype-by-environment interactions (GEI), which is essential for selecting stable and high-performing genotypes under variable production conditions. However, incorporating GEI into GP models increases the dimensionality and computational complexity, risking complex models that are impractical to use on commercial breeding-scale data sets because of run times and computational demands.

This study addresses two primary aims. Firstly, we evaluate the practical benefits of GEI-informed GP for predicting economically important cotton traits. Second, advanced statistical modelling strategies are developed and assessed for integrating genomic and environmental data at scale. We propose a dimensionality reduction approach that combines linkage disequilibrium network analysis with principal component techniques to reduce redundancy while preserving informative variation. Using this reduced dataset, we implement Bayesian linear regression models and, for comparison, deep residual neural networks for genomic prediction. Analyses were conducted on a large multi-environment dataset from the CSIRO cotton breeding program, comprising 3,236 breeding lines, 54 environmental covariates, and 8,049 yield and fibre quality phenotype records collected over 10 years and 9 locations representing 41 year-location combinations. Results demonstrate that generally Bayesian linear regression approaches outperform BG-BLUP models, with all three linear/linear mixed methods providing clearly more reliable performance than the deep learning models. These findings highlight the value of using interpretable statistical models for integrating genomic and environmental information to support selection decisions under diverse environmental conditions.

## Introduction

Cotton, predominantly the tetraploid species *Gossypium hirsutum*, is cultivated in over 70 countries worldwide. It is the leading renewable source of textile fibre (Paterson et al. 2012) and also provides valuable plant-based oil and protein for food and feed (Jabran et al. 2019). The global economic importance of cotton is significantly driven by fibre yield and quality, which are influenced by genetic architecture and environmental conditions. Environmental factors such as temperature and water availability play an important role in shaping cotton performance. Understanding their interactions with genetic variation is essential for developing strategies to breed elite commercial cotton varieties with improved fibre yield and quality under diverse and changing growing conditions.

Conventional phenotype-based breeding methods, while historically effective, are often constrained by their reliance on time-consuming and resource-intensive field trials (Bernardo 2008; Cobb et al. 2019). Most economically important traits are complex and controlled by multiple genes with small effects (Cooper et al. 2009; Hajheidari et al. 2024), making them difficult to improve through marker-assisted selection, and traditional approaches such as phenotype-based selection have limitations. This is because phenotypic expression is often confounded by environmental factors, which can obscure true genetic differences among genotypes and bias the assessment of genetic potential.

Relatively recently, genomic prediction (GP) and/or selection (Meuwissen et al. 2001) have emerged as powerful tools in plant breeding, enabling earlier and more efficient selection of superior genotypes (Crossa et al. 2017; Alemu et al. 2024). Unlike traditional breeding methods that rely heavily on phenotypic evaluations conducted at later growth stages, GP leverages genome-wide molecular marker data to predict genomic estimated breeding values (GEBV) of individuals at an early stage, using statistics and/or machine learning predictive methods (Crossa et al. 2025; Zhang et al. 2025). This approach has the potential to significantly reduce the time and cost associated with field trials while accelerating genetic gain per unit time (Jannink et al. 2010; Voss-Fels et al. 2019). By incorporating genome-wide marker information, GP captures the effects of both large- and small-effect genetic loci, making it particularly effective for complex traits controlled by small-effect genes (Meuwissen et al. 2001; Zhang et al. 2021).

Quantitative traits are shaped not only by genetic factors but also by environmental influences and their complex interactions. This makes their accurate prediction and selection particularly challenging (Doerge 2002; Crossa et al. 2022). In practical breeding programs, the performance of a genotype depends not only on its genetic potential but also on the environment in which it is grown and, critically, on how its genes interact with environmental conditions. Explicitly modelling genotype-by-environment interactions (GEI) (Napier et al. 2023) within genomic prediction frameworks can enable the identification of genotypes with stable performance across diverse environments (Carvalho et al. 2024) or those specifically adapted to target conditions (Halpin-McCormick et al. 2025), thereby enhancing the effectiveness of selection strategies. Moreover, incorporating GEI might also be valuable in the context of climate change, where environmental conditions are becoming increasingly variable and unpredictable (Xiong et al. 2022).

Various multiple-environment GP models have been proposed to account for GEI, either incorporating environmental covariates such as temperature and rainfall or by using location and year as categorical variables. Classical methods such as genomic best linear unbiased prediction (G-BLUP) have been extended to model GEI by additional random-effect terms with covariance structures that combine genomic and environmental components, allowing genotype-specific responses to environmental gradients, forming what is known as a reaction norm model (Jarquín et al. 2014). Factor analytic models (Meyer 2009) take a different approach, using a low-rank structure to efficiently model genetic correlations across environments (Tolhurst et al. 2022). Bayesian methods such as Bayes alphabet (e.g. Bayes A, B and C; Meuwissen et al. 2001; Habier et al. 2011) and their variants allow flexible modelling of marker effects with different prior assumptions and can be extended to incorporate environment-specific effects to represent GEI. More recently, deep learning models (Montesinos-López et al. 2021), including convolutional neural networks (Washburn et al. 2021) and multi-layer perceptron (Yao et al. 2025), have shown good potential in modelling GEI by integrating genomic and environmental data in hierarchical modules (Montesinos-López et al. 2023). These approaches can capture complex, non-additive interactions (Wang et al. 2023), making them attractive for predicting trait performance in untested environments.

Among these GP methods, Bayesian linear regression offers the advantage of not only estimating GEBVs but also providing interpretable effect sizes for individual genetic variants, environmental covariates, and their pairwise interactions (Meher et al. 2022). This explicit modelling of GEI can enhance biological insight and support decision-making in breeding programs. However, incorporating pairwise G×E terms substantially increase the number of parameters, especially with high-dimensional genomic and environmental data, leading to significant computational challenges. Similarly, deep learning approaches, while powerful in capturing complex non-linear interactions, are also computationally intensive, especially when high dimensional data are presented (Montesinos-López et al. 2021; Lourenço et al. 2024). The substantial computational requirements of both G × E Bayesian and deep learning models present barriers to their implementation in routine breeding pipelines, compared with simpler genetic-only models. Efficient strategies for dimensional reduction (Heilmann et al. 2025) are therefore essential for practical applications of these advanced methods in plant breeding. Existing dimensional reduction studies in GP (e.g. Manthena et al. 2022; Song et al. 2025) focused on the genetic data, not GEI terms.

There have been several studies applying genomic prediction in cotton (e.g. Gapare et al. 2018; Islam 2020; Li et al. 2022; Tolhurst et al. 2022); however, most have focused on single-environment scenarios or limited environmental variation. While a few have incorporated multi-environment trial data (Gapare et al. 2018; Tolhurst et al. 2022), efforts to evaluate or develop sophisticated statistical models to account for GEI in breeding scale cotton data remain underdeveloped. As a result, the potential of genomic prediction to fully exploit GEI in cotton breeding has yet to be realised.

Using large-scale data collected from the CSIRO cotton breeding program, spanning multiple years, locations and a wide range of environmental conditions, this study aimed to address two objectives: First, we aim to advance predictive breeding research in cotton by conducting a comprehensive multi-environment GP analysis to address GEI for fibre quality traits including—fibre length, strength, short fibre index, elongation, micronaire, and uniformity— along with lint percentage and lint yield. Second, we aim to develop an efficient dimensionality reduction approach that combines linkage disequilibrium (LD) network analysis (Li et al. 2018) with principal component techniques, addressing the challenge of high-dimensional G×E data. This approach is designed to reduce redundancy in genomic and environmental features while preserving informative variation. The reduced dataset is then used for genomic prediction employing either Bayesian linear regression models (Habier et al. 2011; Li et al. 2024) or deep learning architectures (He et al. 2016), enabling robust and scalable prediction across diverse environments. The importance of this study is because both the accurate modelling of GEI and the development of GP models that can be deployed in commercial plant breeding scenarios are vital for the continued improvement of crop varieties.

## Results

To evaluate the performance of genomic prediction across multiple environments, we integrated multi-environment phenotypic records with genome-wide SNP data to model genetic main effects and genotype-by-environment interactions (GEI). Linkage disequilibrium (LD) structure was characterised and exploited to reduce the dimensionality of GEI terms prior to prediction. We implemented and compared four genomic prediction models, including Bayesian genomic best unbiased linear predictor (BG-BLUP), Bayes C, LD-Bayes and Multi-module deep learning method based on residual network architecture (ResNet), to assess their relative accuracy across fibre quality and yield traits.

### Phenotypic variation and genomic heritability of fibre quality and yield traits

Across multiple environments, substantial phenotype variation was observed for all traits of interest (Table 1). Coefficients of variation indicated higher dispersion for LY, EL and SFI. The genomic heritability, defined as the portion of the phenotype variance explained by the SNP data varied among traits. LEN and STR were the most heritable 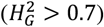, followed by 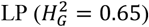. In contrast, LY exhibited the lowest heritability 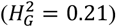, with a substantial proportion of its variation attributable to G×E interactions 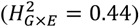 Fibre quality traits, UNI, SFI, EL and MIC, were moderately heritable 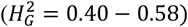.

**Table 1.** Phenotypic variation and genomic heritability (*h*^2^) for eight traits across all environments.

| Trait | Unit | Mean | Range | SD | CoV | $H_G^2$<br>(SD) | $H_{G \times E}^2$<br>(SD) |
| --- | --- | --- | --- | --- | --- | --- | --- |
| LEN | mm | 1.25 | 1.07-1.46 | 0.05 | 4.0 | 0.75<br>(0.02) | 0.07<br>(0.01) |
| UNI | % | 84.17 | 80.2-88.4 | 1.17 | 1.4 | 0.40<br>(0.02) | 0.18<br>(0.01) |
| SFI | % | 5.51 | 2.3-9.9 | 1.30 | 23.6 | 0.43<br>(0.02) | 0.17<br>(0.01) |
| STR | g tex<sup>-1</sup> | 31.83 | 26.8- 39.3 | 1.45 | 4.6 | 0.71<br>(0.02) | 0.09<br>(0.01) |
| EL | % | 5.60 | 1.9-8.7 | 1.21 | 21.6 | 0.40<br>(0.02) | 0.29<br>(0.02) |
| MIC | — | 4.22 | 2.6-5.5 | 0.40 | 9.5 | 0.58<br>(0.02) | 0.13<br>(0.02) |
| LY | kg ha<sup>-1</sup> | 2632.83 | 436.4-4420.8 | 497.94 | 18.9 | 0.21<br>(0.02) | 0.44<br>(0.02) |
| LP | % | 43.01 | 32.2-50.4 | 2.5 | 5.8 | 0.65<br>(0.02) | 0.17<br>(0.01) |

SD=Standard deviation,
CoV= Coefficients of variation = $\frac{SD}{MEAN} \times 100$ .
$H_G^2$ =genomic heritability=proportion of phenotype variance explained by the SNP data
$H_{G \times E}^2$ =proportion of phenotype variance explained by the interaction between SNPs and environments

### Genotyping and linkage disequilibrium analysis

The DArT genotyping array initially contained 8,707 SNPs. After filtering SNPs with MAF<2.5%, heterozygosity rate>80%, and missing genotype rates>20%, 7,532 informative SNPs distributed across 26 chromosomes (A1-A13; D1-D13) of cotton were retained for subsequent LD analysis and genomic prediction.

LD network analysis classified these SNPs into 2,072 LD blocks, with the number of blocks per chromosome ranging from 46 to 139 (Table S1; Figures S1). On average, each block comprised 3.6 SNPs, although the distribution was uneven: for example, 13 blocks each comprising over 50 SNPs and 1,239 blocks only represented by a single SNP. Chromosome-wise, LD results revealed extended haplotype blocks in mid-chromosomal regions and finer blocks toward distal arms across both A and D sub-genomes (Figure S1). While the size of some blocks is quite large, the LD block pattern across chromosomes is generally in line with the distribution of protein-coding genes from which the SNPs were mainly derived.

### Dimensional reduction of GEIs

Model-based clustering partitioned the ECs into nine distinct groups. PCA was then separately applied to each of the resulting 2,072 × 9 (LD-block × EC-group) GEI matrices, producing 51,035 principal components, representing a major reduction from the initial 7,532 (SNPs) × 54 (environmental covariates) (406,728) GEI terms.

### Genomic prediction

Across all traits, the BG-BLUP, Bayes C, and LD-Bayes methods yielded comparable levels of prediction accuracy when averaged over the CV1 runs, and all three outperformed the ResNet approach (Figure 1; Table S2). Among the genomic prediction models, Bayes C showed the strongest overall performance, achieving the highest mean accuracy for three (LEN, EL, and LP) of the eight traits, and performed similarly to BG-BLUP or LD-Bayes for the remaining traits.

**Figure 1:**
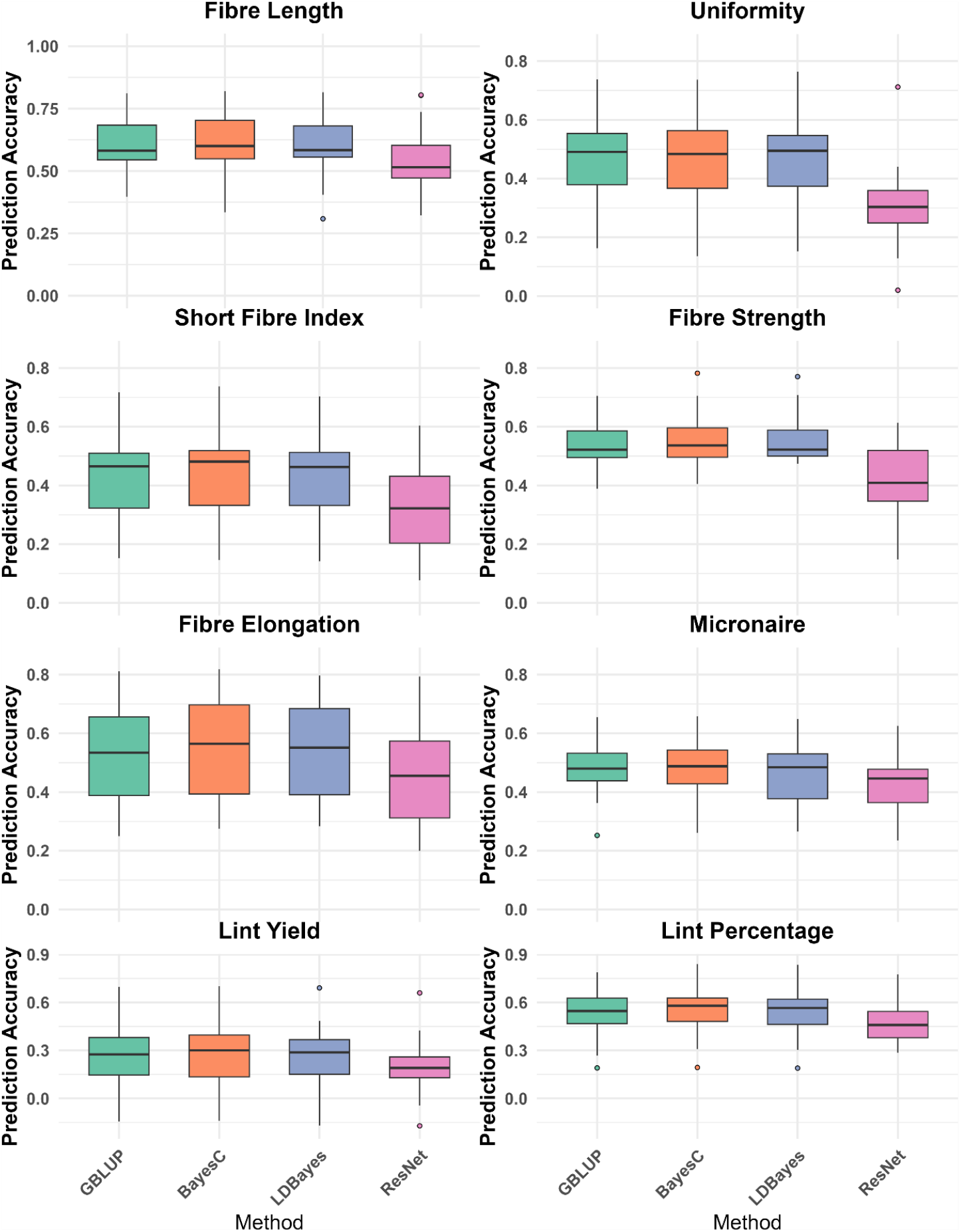
Distribution of prediction accuracies obtained across leave-one-environment-out cross-validation runs for eight cotton traits including Fibre length, Uniformity, Short fibre index, Fibre Strength, Elongation, Micronaire, Lint yield and Lint percentage using four genomic prediction methods (Bayesian genomic best linear unbiased predictor or BG-BLUP, Bayes C, LD-Bayes, and Residual network or ResNet). Each panel corresponds to a trait and displays boxplots summarizing the variability and central tendency of accuracy for each method.

Using the Bayes C method, which overall provided the highest prediction accuracy among the methods, as an example, we further compared the performance of G×E model (Equation 2), with two reduced models: the G+E model, which captures only the additive effects of genotypes and environments while ignoring the GEI, and the G model, which includes genotype effects only. The results (Figure 2; Table S3), based on distributional and average prediction accuracies across multiple cross-validation replicates, show that the G+E model achieved an increase in prediction accuracy by 13 to 96% compared with the G model across all traits (LEN: 0.14, UNI: 0.20, SFI: 0.25, STR: 0.22, EL: 0.96, MIC: 0.28, LY: 0.29, LP: 0.23). The full model with the additional inclusion of GEI effects resulted in further improvements of prediction accuracy relative to the G+E model, though the extent varied with the traits (2–23%; LEN: 0.07, UNI: 0.12, SFI: 0.13, STR: 0.10, EL: 0.08, MIC: 0.02, LY: 0.23, LP: 0.15).

**Figure 2:**
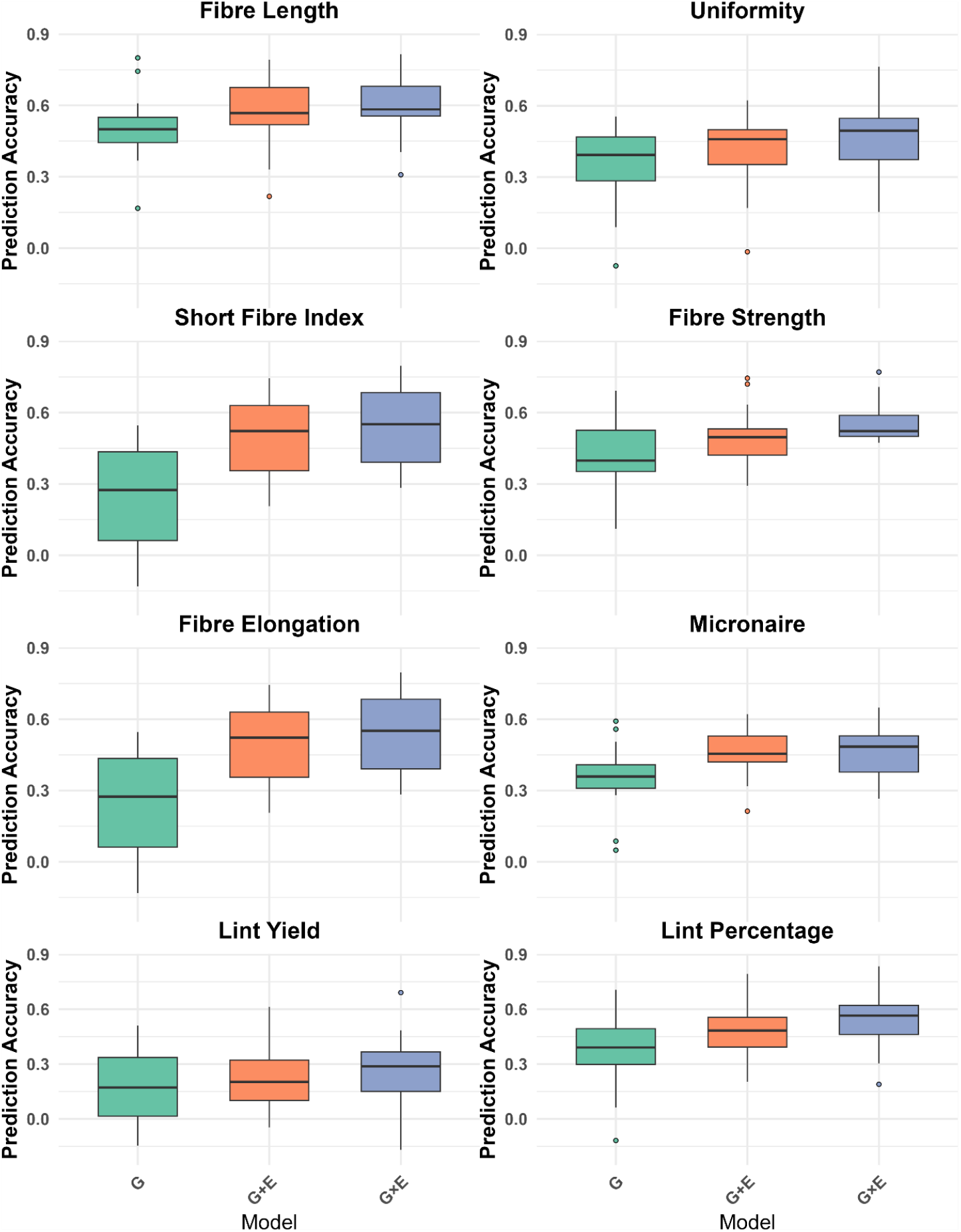
Distribution of prediction accuracies obtained across leave-one-environment-out cross-validation runs for eight cotton traits including Fibre length, Uniformity, Short fibre index, Fibre Strength, Elongation, Micronaire, Lint yield and Lint percentage using the Bayes C method. The three model under comparison are (i) the full G×E model which includes additive genetic effects, environmental effects and their interactions, and (ii) the G+E model, which only includes the additive effects of genetics and environments, and (iii) the G model which only includes the genetic effects. Each panel corresponds to a trait and displays boxplots summarizing the variability and central tendency of accuracy for each method.

Prediction accuracies were correlated with the genetic architecture of the traits. Across the eight traits, the prediction accuracies obtained using Bayes C showed a strong positive correlation with the square root of the genomic heritability (r=0.88), indicating that traits with higher heritability consistently yielded higher predictive power. Specifically, STR and LEN with the highest 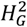 values (Table 1) also achieved the highest accuracies (Figure 1, 2; Table S2, S3), whereas the lowest accuracies were observed for LY which also has the lowest genomic heritability. The same tendency was observed for other methods (Results not shown).

## Discussion

Our work makes two major contributions to GP. First, it presents a comprehensive multi-environment genomic prediction study on a major crop, upland cotton, using breeding-scale datasets. Second, it introduces a linkage-disequilibrium–based dimensionality-reduction procedure for modelling genotype-by-environment interactions (GEI). This approach substantially reduces data dimensionality and makes computation feasible for a broad range of models, including Bayesian linear regression, neural networks, and deep learning methods.

### Partitioning genetic and GEI effects

Unlike conventional multi-environment field trials that rely solely on phenotypic observations across replicates and environments, our study integrated genome-wide marker data with environment covariates (EC) derived from weather records. This allowed us to partition phenotypic variance into components attributable to pure genetic effects (i.e. genomic heritability) 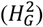 and GEI effects 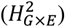, conditional on observed environmental effects. Our estimates of genomic heritability 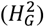 across eight traits align closely with those reported by Li et al. (2022), maintaining a strong correlation (*i*.*e*. Pearson correlation = 0.84) between the two studies. Notably, the estimated 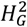 values in this work were generally higher, likely reflecting the increased statistical power from a larger sample size compared to the pre-2018, single-site (MV) population used previously in Li et al. (2022). A key advancement of this work is that we move beyond these pure genetic effects to explicitly account for GEI. The results showed that lint yield was more strongly influenced by GEI than by main genetic effects, whereas lint percentage was predominantly controlled by genetic effects. Among the six fibre quality traits analysed, fibre length and fibre strength were strongly genetically determined traits. These findings are consistent with previous studies (Liu et al. 2011; Virk et al. 2023; Snider et al. 2013; Scarpin et al. 2025). For example, Virk et al. (2023) reported that lint yield is more strongly influenced by environmental effects than by genotype, while lint percentage, fibre length, and fibre strength are primarily under genetic control based on 19 years of multi-location trials in Arkansas.

### Comparison between different GP methods

In this study, we compared four GP methods that differ in their underlying assumptions and strengths, allowing us to evaluate models’ performance across a range of traits. Among the linear models, BG-BLUP assumes marker effects follow a common normal distribution with a shared variance, making it most suitable for highly polygenic traits where all markers contribute small additive effects (Meuwissen et al. 2001; VanRaden 2008). In contrast, Bayes C (Habier et al. 2011) and its variant LD-Bayes (Li et al. 2024) assign marker-specific prior distributions, enabling them to accommodate genetic architectures where a small number of loci have moderate-to-large effects and the remaining markers contribute little. Accordingly, these Bayesian linear regression methods are generally more effective for oligogenic or sparse-effect traits, while BG-BLUP performs best for traits governed by many small-effect loci. We additionally included ResNet (He et al. 2016; Montesinos-López et al. 2023) as a non-linear deep learning method, which we hypothesised might be able to capture non-additive genetic patterns such as dominance and epistasis effects that linear models cannot implicitly represent (Montesinos-López et al. 2024).

Overall, the performance of BG-BLUP, Bayes C and LD-Bayes was similar (Figure 1; Table S2). However, the higher accuracies obtained for STR and LP with the latter two methods suggest that these traits may have a less polygenic architecture, potentially influenced by a smaller number of loci with moderate-to-large effects (Fang et al. 2021; Chen et al. 2022; Niu et al. 2022).

The genomic prediction results showed that the ResNet model generally performed worse than the other three methods (Figure 1; Table S2). This pattern suggests that strong dominance or epistatic effects—which deep learning models are often capable of capturing— may not be prominent in the current dataset. It is also important to note that the performance of ResNet, like many other deep learning approaches, is highly sensitive to the choice of network architecture, hyperparameters, and initialization settings. Further optimization of these aspects may improve its predictive performance (Montesinos-López et al. 2024), although such model-tuning efforts are beyond the scope of the present study.

Another important difference among the four methods concerns the form of the input data. BG-BLUP uses the genomic relationship matrix G and the GEI covariance matrix GE (*n* × *n* matrices; *n* is the number of phenotype records), both of which can be pre-computed from genotype (VanRaden 2008) and environment data (Costa-Neto et al. 2021), respectively. In contrast, Bayes C, LD-Bayes, and ResNet all use SNP genotypes directly, together with the pairwise GEI terms, as input variables. Consequently, these three methods benefit from the LD-based dimensional reduction applied to the GEI terms, which substantially reduces the number of predictors entering their models. From a computational perspective, BG-BLUP has a non-linear (approximately cubic) increase in computational complexity with respect to the number of phenotype records (*n*), because mixed model equations require operations such as matrix inversion or decomposition of an *n* × *n* covariance matrix. By contrast, the computational cost for Bayes C, LD-Bayes, and ResNet increases approximately linearly with *n*. Their computational burden is instead dominated by the number of regression variables (SNPs and GEI terms). With the LD-based dimensional reduction applied to the GEI inputs, the number of predictors is greatly reduced, making the computational requirements of these methods far less limiting.

### Comparison of G, G+E, and G×E models

For fibre quality traits including LEN, EL, MIC, SFI and UNI, incorporating environmental main effects (G+E) substantially improved prediction accuracy relative to genomic effects (G) alone, indicating strong and predictable environmental influences. However, further inclusion of GEI terms resulted in marginal additional gains. This suggests that environmental responses for these traits are largely consistent across genotypes, with limited genotype-specific deviations, likely reflecting the close genetic relatedness and reduced allelic diversity of the breeding lines used in this study.

Interestingly, the inclusion of GEI effects improved genomic prediction accuracy of LY by a comparable altitude for traits like LP (Figure 2; Table S3), despite lint yield exhibiting a much larger overall G×E variance component than LP (Table 1). This indicates the G×E effects associated with LP are more consistently captured by the model, reflecting more structured and environment-specific patterns of genotype response affecting fibre–seed partitioning. In contrast, although lint yield shows a large G×E component, much of this interaction variance might be poorly predictable, limiting the gain from incorporating GEI into prediction models. This pattern is consistent with previous QTL and/or environment studies reporting stable genetic control for LP (Shang et al. 2016; Niu et al. 2022; Baghyalakshmi et al. 2024) but highly environment-driven and unstable performance for lint yield (Sharif et al. 2025).

### LD block patterns

Segmenting chromosomes into LD blocks based on pairwise correlation of SNPs is a common practice to compress high dimensional genotype data to lower-dimensional representations, while preserving genetic patterns. This practice has been adopted in mapping quantitative trait loci, genome-wide association studies, and GP of both animals and plants (e.g. Li et al. 2018; Ye et al. 2022; Li et al. 2024). In this study, we successfully conducted LD clustering for dimensional reduction to mitigate the computational challenges imposed by incorporating GEI in GP. The LD block sizes of the cotton lines used in this study varied significantly, from 0.32 kb to 104 Mbp (for those blocks with more than 1 SNP), with the small and large LD blocks mainly located at the distal regions and around centromeres of chromosomes, respectively, particularly in A sub-genome (Figures S1). This distribution of LD blocks mirrors the known recombination landscape in cotton—suppressed around centromeres and elevated distally—and is consistent with the higher recombination per megabase reported for the D sub-genome relative to A sub-genome (Shen et al. 2017). Some of the large LD blocks could be attributable to the nature of low genetic diversity of the breeding lines used in this study. Intense artificial selection for high fibre yield and quality among highly genetically related materials tends to create long and stable haplotype blocks. This is consistent with the notion of slower LD decay rate in modern upland cotton cultivars (lower genetic diversity) compared to wild/landrace upland cotton accessions (higher genetic diversity) (Wang et al. 2017). The LD blocks reported here were generated based on relatively small number of SNPs (7,532). Continuing reductions in sequencing costs would make it feasible to have more SNPs by sequencing all materials used in GP, and therefore it is of interest to understand the impact of increased number of SNPs on LD block patterns and, in turn, the impact of LD block patterns on GP accuracy. Once SNP density becomes sufficiently high, LD block patterns are expected to converge, at which point further improvements in prediction accuracy would likely depend on population composition rather than refinements in LD structure.

In conclusion, this study presents a genomic prediction analysis using a large breeding-scale multi-environment cotton dataset, demonstrating that explicitly modelling GEI effects can improve prediction accuracy for lint yield and most fibre quality traits. By applying a linkage-disequilibrium-based dimensionality reduction approach, we condensed high-dimensional GEI features into a tractable form, enabling computational feasibility of subsequent Bayesian regression and deep learning analyses. As an extension of earlier findings (e.g. Li et al. 2022), these results reinforce that genomic prediction can achieve prediction accuracies applicable in commercial cotton breeding programs and incorporating environmental covariates and GEI can considerably improve model predictive performance, providing further evidence that such approaches can further empower breeders to effectively and efficiently use the resources in their breeding activities and deliver impactful new varieties for diverse production environments.

We also recognize several limitations and opportunities. First, although we incorporated a rich set of environmental covariates, continued expansion to include additional information such as remote-sensing indices (Fritsche-Neto et al.2025) may capture additional variance components and further stabilize predictions in highly variable environments. Second, because no single model was uniformly superior, routine benchmarking and model ensembles might be useful to ensure robust predictive performance. Finally, the question remains whether extending the framework to multi-trait prediction with explicit genetic and environmental covariance structures may enhance accuracy and biological interpretability for correlated traits.

## Materials and Methods

### Phenotypic data

The traits analysed in this study include lint yield (LY; kg ha^-1^), lint percent (LP; percentage of lint out of harvest seed cotton, %), and six fibre quality traits: fibre length (LEN; upper half mean length of sample), uniformity (UNI; ratio of mean fibre length to upper half-mean length, expressed as %), short fibre index (SFI; proportion by weight of fibre shorter than 12.7 mm), strength (STR; force required at the breaking point for a bundle of fibres of a given weight and fineness, g tex^-1^), elongation (EL; extension ability of a bundle of fibres up to its breaking point, expressed as a % increase over its original length), and micronaire (MIC, a composite measure of fibre fineness and maturity based on air permeability, unitless).

All phenotype data were collected from experiments conducted under fully irrigated conditions, spanning over 10 seasons (2014–2023) at 9 locations within conventional cotton growth areas in Eastern Australia (Table S4), though not all locations were represented in every year. As described in Li et al. (2022), experiments at each site–year combination used a row– column design with four replicates generated by CycDesigN software (VSN International, UK). Each plot comprised three cotton rows (10–12 m long, 1 m spacing) at a planting density of approximately 10–12 plants m^−2^. All trials followed commercial management practices, including irrigation and weed/pest control (Cotton Research and Development Corporation, 2025). Furrow irrigation was applied when required, approximately every 10–14 days (≈1 ML ha^−1^ per irrigation) from December to March. At ~60% open bolls, crops were defoliated with thidiazuron, and unopened bolls were treated with ethephon, followed by a second application 7–10 days later.

At harvest, seed cotton was mechanically harvested from the middle row of each plot with a spindle picker (modified Case International 1822) and weighed. The outside rows were not harvested and acted as buffers to minimise the edge and neighbouring competition. LP was determined by ginning a 300 g sub-sample of seed cotton from plot harvests using a 20-saw gin with a pre-cleaner (Continental Eagle, Prattville, AL U.S.A.) and was subsequently used to calculate lint yield (kg ha^−1^). Lint samples were collected and tested for fibre quality using a Spinlab High Volume Instrument (HVI) model 1000 (Uster Technologies AG, Uster, Switzerland).

Individual experiment’s phenotype data were analysed using linear mixed models taking account of two-dimensional spatial variation, as described in Liu et al. (2015) and Li et al. (2022). The best linear unbiased estimates of test lines from individual trial analysis were pooled and used as the phenotype value in the genomic prediction analyses.

In total, 3,236 conventional cotton breeding lines were phenotyped through the breeding program and used for subsequent genomic analyses. These lines were predominantly at the F_4_ – F_6_ generation, depending on the initial selfing generation of breeding families used to grow into the populations for single-plant selection to derive fixed lines, which were subsequently evaluated and progressed or dropped stage-by-stage based on performance data from the trials conducted in multiple location and year (Liu and Constable, 2017). The dataset included 3,236 released cultivars and breeding lines undergoing different stages of testing. Of these, 937 lines (phenotyped before 2018) were previously analysed by Li et al. (2022) based on genotyping data generated using the CottonSNP63K array (Hulse-Kemp et al. 2015). In this study, they were re-genotyped using a DArT sequencing platform (see Genotyping and SNP Filtering) with a full set of phenotype data from the breeding base station, Myall Vale, as well as the other farm locations (Table S4). In total, 8,049 phenotypic records were available across all traits, corresponding to an average of 2.5 observations per line. Of the 3,236 lines, 2,010 were evaluated in more than one environment, in terms of year and location combinations.

### Environment data collection and phenology modelling

Climate or environmental data were obtained in two different ways: (i) from weather stations located at the experiment site; (ii) from the SILO climate database (Jeffrey et al. 2001), where weather station data were not available.

The measurable climate data consisted of daily records for solar radiation, maximum and minimum temperature, rainfall, and relative humidity at maximum temperature. Vapour pressure deficit (VPD) was also calculated on the basis of temperature and humidity (Conaty et al. 2014). The environmental covariate records were available for each day of the growing season from planting to harvest.

Six phenological growth-stages were defined as the key stages of crop development: first emergence, first square, first flowering, peak flowering, open boll and crop maturity. Growth-stage development in cotton can be approximated as a mathematical function of day-degrees (Constable and Shaw, 1988). The day-degree is a thermal-time metric intended to reflect the potential rate of plant development based on the temperature range for a given day. Day-degrees (DD) were calculated using the following phenology model (Bange et al., 2022):

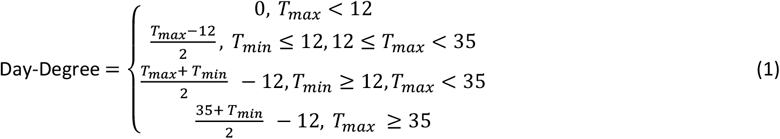

where T_max_ and T_min_ are the maximum and minimum temperatures in Celsius recorded for a given day. The day-degree targets assumed to coincide with the phenological growth-stages of cotton were given in Table S5. Note that, when *T*_min_<11 °C occurs before first square or first flower, the event can delay cotton growth and development, requiring an additional 5.2 DD to overcome this setback. On the basis of the formula (1) and the day-degree targets commonly used in the industry (Table S5), the start and end dates of the six growth periods were estimated for test environments, when referring to the recorded sowing dates. Within each growth stage, the environmental covariates were defined as (i) mean of daily T_max_, (ii) mean of daily T_min_, (iii) mean of daily average temperature (T_max_ + T_min_)/2, (iv) number of hot days (i.e >35°C), (v) number of cold days (i.e <12°C), (vi) cumulative daily precipitation, (vii) number of rainy days, (viii) cumulative daily radiation, and (ix) mean of VPD at T_max_.

### Genotyping and SNP filtering

Leaf samples from the 3,236 lines were collected across multiple seasons. After freeze-drying, samples were sent to Diversity Arrays Pty Ltd. (DArT, Canberra, Australia) for DNA extraction and genotyping using proprietary protocols. Genotyping was performed using DArTag™, a genotyping-by-sequencing platform based on a custom array of 8,707 single nucleotide polymorphism (SNP) markers. These markers were selected from whole-genome resequencing data of diverse Australian cotton varieties and breeding lines, many of which contributed to the publicly available CottonSNP63K array (Hulse-Kemp et al., 2015). SNP calling followed DArT standard procedures, and markers were mapped to the upland cotton reference genome CSX8308 (Sreedasyam et al., 2024). Quality control involved filtering SNPs with >20% missing data, minor allele frequency <2.5%, and heterozygosity rate>80%. Missing genotypes of the remaining SNPs were imputed using missForest (https://github.com/stekhoven/missForest), a non-parametric algorithm that employs Random Forests to iteratively predict missing values, effectively handling complex, non-linear relationships without strong distributional assumptions (Stekhoven and Bühlmann 2012; You et al. 2023).

### Estimation of linkage disequilibrium blocks

Physically adjacent SNPs are often in linkage disequilibrium (LD), forming correlated genomic regions. To capture this structure, genome-wide SNPs were classified into LD blocks, which were later used to guide the dimensional reduction of GEIs and incorporated as prior information in Bayesian regression models for genomic prediction. LD blocks were derived using the LD network clustering (LDn-clustering) algorithm (Li et al. 2018). Briefly, each chromosome was divided into approximately equal-sized and non-overlapped windows, and the pairwise LD (measured as *r*^2^) among SNPs within each window was calculated. Based on these pairwise LD values, SNPs with high LD were grouped into blocks using a network-based clustering procedure (Li et al. 2018). The method was implemented using the function “LDnClustering” in the R package LDna (Kemppainen et al. 2015; https://github.com/petrikemppainen/LDna).

### Estimation of environmental covariates groups

A classical model-based clustering approach named Gaussian mixture (Fraley et al. 2012) was used to cluster environmental covariates (EC) into environment groups. An optimal number of clusters was determined by Bayesian information criterion.

### Linear regression model for multi-environment data

The linear regression model to analyse multi-environment data is presented as:

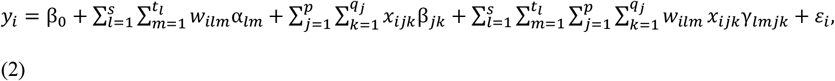

where, *y*_*i*_ is the phenotype of the observation *i* (*i*=1,…, *n*), *w*_*ilm*_ is the value of the EC*m* (*m*=1,…,*t*_*l*_) in group *l* (*l*=1,…, *s*) at the environment (e.g. a unique combination of year and field trial site) where the observation *i* is taken, and *x*_*ijk*_ is the genotype value of SNP *k* (*k*=1,…,*q*_*j*_) located within the LD block *j* (*j*=1,…,*p*) of the line *i*, coded as *x*_*ijk*_=-1,0, 1 for genotypes AA, AB and BB, respectively, *β*_0_ is the fixed intercept term representing the population mean, *α*_*lm*_ is the effect of the environmental variable *m* in the environmental group *l*, β_*jk*_ is the main additive genetic effect of SNP *k* in the LD block *j, γ*_*lmjk*_ is genotype-by-environmental interaction (GEI) effect, i.e., the interaction between EC *m* in the environment group *l* and the SNP *k* in the LD block *j, ε*_*i*_ is the residual which is mutually independent and follows a normal distribution 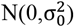, with unknown variance 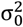.

Model (2) can be specified as a likelihood function:

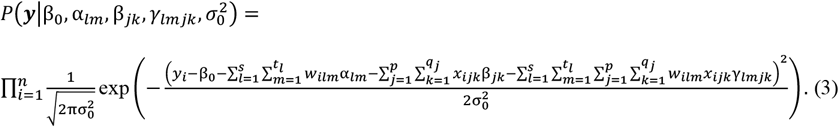

### Dimensional reduction of Genotype × Environment interactions

In equation (3), the high dimensionality of the GEI terms

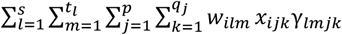

substantially increases memory and computational requirements in genomic evaluation. To reduce the computational cost, the following dimensional reduction strategy was applied. Firstly, the interaction terms *w*_*ilm*_*x*_*ijk*_ were clustered according to environment group *l* and genomic region *j*, forming interaction clusters defined as Group_*l*×*j*_=Group_*l*_ × Group_*j*_. Secondly, within each interaction cluster Group_*l*×*j*_ (*l*=1, …, *s*; *j*=1, …, *p*), principal component analysis (PCA) was applied to the corresponding matrix of GEI terms 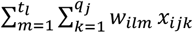. The leading PCs, denoted by z_iljg_ (*g*=1, …, *r*_*l*×*j*_) which together explained at least 70% of the total variation, were retained and used as a replacement for the original GEI terms. Since environmental covariates within each environment group and SNPs within each LD block are expected to be highly correlated, only a small number of PCs should be sufficient to capture the majority of the variation in the original GEI data (i.e. *r*_*lj*_ << *t*_*l*_ × *q*_*j*_). Consequently, this PCA-based strategy substantially reduces the model dimensionality, and lowers the computational cost of the subsequent GP analysis.

Accordingly, equation (3) can be reformulated as

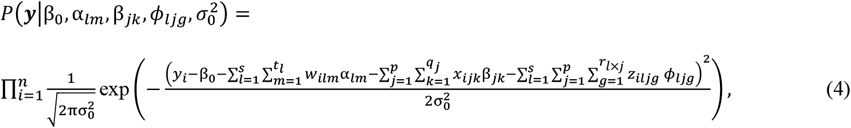

where *z*_*iljg*_ denotes the score of the *g*th PC in the interaction cluster Group_*l*×*j*_ for individual *i*, and *ϕ*_*ljg*_ is the corresponding PC effect. All remaining symbols in equation (4) are defined identically in equation (3).

### Bayes C

The model parameters in the likelihood (4) are assigned prior distributions. Combining the likelihood with these priors yields the joint posterior distribution

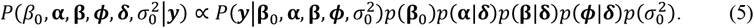

We adopt a spike-and-slab regression framework, commonly referred to as Bayes C in the genomic selection literature (Habier et al., 2011). The intercept term *β*_0_ is assigned a non-informative uniform prior as *β*_0_~*Uni*(−∞, ∞).

The environmental effect α_*lm*_, additive genetic effect *β*_*jk*_ and GEI effects *ϕ*_*ljg*_ are assigned spike-and-slab priors (Ishwaran and Rao 2005; O’Hara and Sillanpää 2009). Specifically, each effect follows a mixture distribution of a normal distribution and a point mass at zero as:

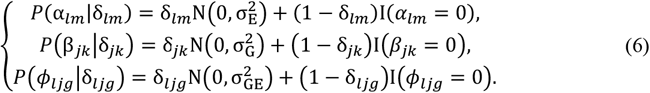

Here, *δ*_lm_, *δ*_jk_, and *δ*_ljg_ are binary indicator variables. When *δ*=1, the corresponding effect is included in the model and assumed to follow a normal distribution with component-specific variance. When *δ*=0, the effect is excluded and fixed at zero. All active environmental, genetic, and GEI effects are assumed to share common variances 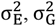 and 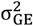, respectively.

In equation (6), the indicator variables are assigned independent Bernoulli priors:

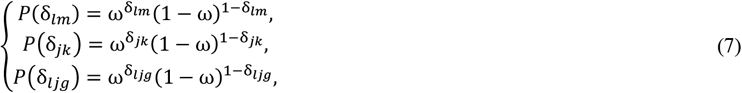

where ω represents the expected proportion of non-zero regression coefficients. A Beta hyperprior is further assigned to ω :

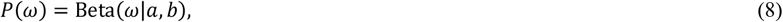

with *a*=*b*=25, yielding a prior mean of *a*/(*a*+*b*) = 0.5, yielding a prior mean of 0.5 and a symmetric but informative prior that favours moderate inclusion rates, reflecting no directional preference toward either inclusion or exclusion of regression covariates.

The residual variance 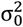 and the component-specific variances 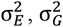 and 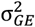 in (4), and (6) are each assigned scaled inverse chi-squared priors:

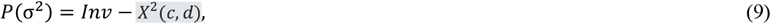

with degrees of freedom c fixed to be 5/2. The scale parameters are specified as

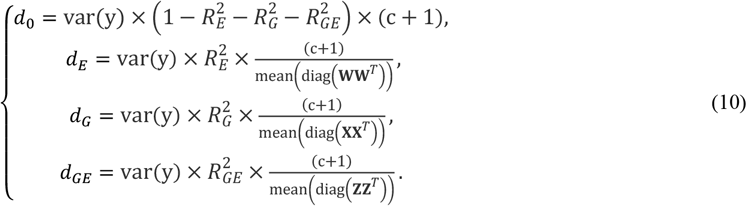

where **W, X** and **Z** denote the design matrices for environmental covariates *w*_*ilm*_, SNPs *x*_*ijk*_ and GEIs *z*_*iljg*_, respectively.

These hyper-parameter settings extend the defaults proposed by Pérez and de los Campos (2014) for Bayes B and C models by explicitly incorporating environmental and GEI effects. The parameters 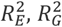, and 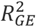 represent prior expectations of the proportions of phenotype variance explained by environment, genetic and GEIs, respectively. In practice, we specify 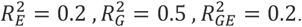.

Given the conditional conjugacy of all priors, posterior inference is conducted using a Markov Chain Monte Carlo (MCMC), or more specifically a Gibbs sampling algorithm. A total of 60,000 MCMC iterations were generated, with the first 10,000 samples discarded as burn-in. The remaining 50,000 samples were thinned by retaining every 50^th^ iteration, resulting in 1,000 posterior samples for inference.

### LD Bayes

The method was originally introduced for quantitative trait locus mapping in a cotton multi-parent advanced generation inter-cross population (Li et al. 2024). The key difference between this approach and Bayes C is that LD Bayes assumes marker effects belonging to different LD blocks share block-specific variance components, rather than a single common variance. Under this framework, ECs in different environment groups, genetic markers in different LD blocks, and GEI terms in different interaction groups are each assigned group-specific variance parameters. Specifically, the spike-and-slab priors are defined as

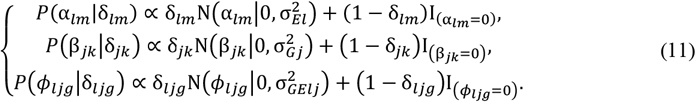

Each group-specific variance parameter is assigned a scaled inverse-chi-squared prior,

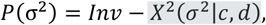

with degree of freedom fixed at c=5/2. The scale parameters are specified as

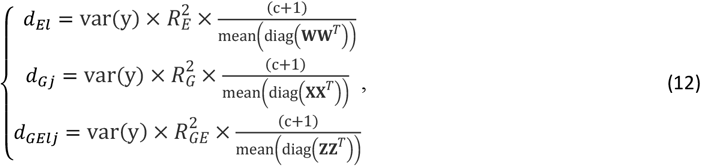

All remaining priors and hyperpriors are specified identically to those used in the Bayes C model.

### Bayesian G-BLUP model

The BG-BLUP model is specified as

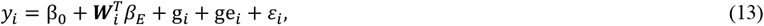

Where 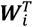 is the vector of the observed ECs where observation *i* is taken, *β*_*E*_ is the corresponding fixed-effect coefficients. The term *g*_*i*_ is the random genetic effect for observation *i*, and *ge*_*i*_ captures the GEI effects. The phenotype data *y*_*i*_, the model intercept *β*_0_, and the residual error *ε*_*i*_ are defined as in Equation (2).

The random effects ***g***=( *g*_1_, …, *g*_n_) and ***ge****=*( *ge*_1_, …, *ge*_n_) follow multivariate normal distributions as:

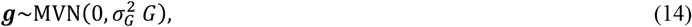

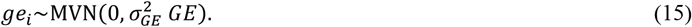

where E represents an environmental kernel measuring the similarity between the environments calculated using the classical variance-covariance matrix of the environment covariates (Costa-Neto et al. 2021), and G is the well-known genomic relationship matrix (vanRaden 2008). In practice, the E matrix and GRM were calculated using R package EnvRtype (Costa-Neto et al. 2021) and rrBLUP (Endelman 2011), respectively. Matrix GE was calculated as the Hadamard product between G and E, as a way to introduce GEI to the G-BLUP model. Under the Bayesian formulation, the variance components 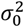 (the variance for *ε*_*i*_),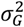 and 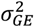 were all assigned Scaled inverse chi-squared prior

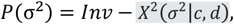

where the degree of freedom *c* was all specified to be 5/2 for these variance components, and the scale parameters are defined as

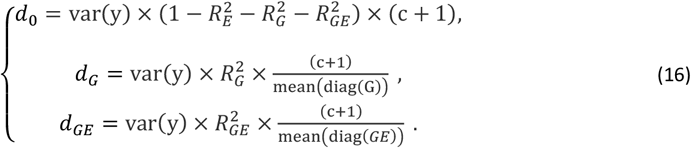

### Genomic heritability

Conditional on observed environmental effects, the proportion of phenotype variance explained by SNP markers, often referred to as genomic heritability (de los Campos et al. 2015), is defined as

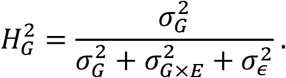

Similarly, the proportion of phenotype variance attributable to the GEI effects is defined as

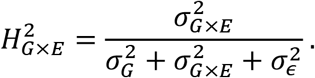

The posterior distributions of the variance components 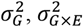, and 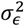 were estimated by MCMC, allowing both point and uncertainty estimates of 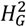 and 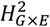 to be derived.

### Multimodal deep learning model

The above described Bayesian approaches all rely on linear and additive assumptions, which may limit their ability to capture the non-additive genetic effects and complex genotype– environment relationships commonly observed in real data. Multimodal deep learning (Ngiam et al. 2011; Montesinos-López et al. 2024) provides a unified and flexible framework capable of modelling such nonlinear and higher-order interactions. To exploit complementary signals contained in genomic markers (i.e. g_*i*_), environmental descriptors (i.e. *e*_*i*_) and their explicit interaction (i.e. g*e*_*i*_), we designed a three-branch encoder–fusion architecture (Figure S2). Each branch is a lightweight stacked residual network, ResNet (He et al. 2016), that converts its modality-specific input into a low-dimensional latent embedding; residual connections mitigate vanishing gradients and enable deeper feature extraction without information loss.

An encoder (Vaswani et al. 2017) is a neural-network module that takes raw data and transforms it into a compact numerical representation that captures key patterns relevant for prediction. In this work, three encoders are used: a genomic encoder (G-ENC), an environmental encoder (E-ENC), and an interaction encoder (G×E-ENC). In G-ENC, genomic marker (i.e. *g*_*i*_) data pass through two residual blocks, each comprising a fully connected (FC) layer, batch normalisation (BN), Rectified Linear Unit (ReLU) activation and dropout (*p* = 0.20) (Goodfellow et al. 2016). The first block has 256 units and the second 128 units, producing a 64-dimensional latent vector *z*_G. As for E-ENC, the ECs are fed into the same residual template with an initial width of 128 units, producing embedding z_E. G×E-ENC encodes explicit GEIs (after dimensional reduction) with a residual path identical to E-ENC, generating embedding vector *z*_GE.

The three embeddings are concatenated and passed to two FC residual blocks (128 and 64 units), followed by a linear output neuron that returns the estimated phenotype values for each target trait. Batch normalization, dropout, and L2 (*λ* = 1×10^-4^) regularisation are applied throughout to prevent over-fitting. The network is optimised with Adam (initial learning rate = 1 × 10^−3^, *β*_1_ = 0.9, *β*_2_ = 0.999) (Kingma and Ba 2014) under mean-squared-error loss. An exponential decay scheduler and early stopping (patience = 15) further stabilise training. The model was trained on a computer with 16 GB NVIDIA RTX 4080 and 64 GB RAM.

### Assessing Model Predictability

We implemented a leave-one-environment-out cross-validation (CV) scheme to evaluate the predictability of different models. In each round, one environment (a year-location combination) was used as the test set, and the remaining environments formed the training set. To avoid information leakage, we excluded any lines present in both sets. For example, when predicting the phenotypic performance of lines collected at location MV in 2020, we excluded data of these lines from all other locations in that year because those locations contained the same lines evaluated at MV. This strategy reflects the scenario of predicting untested lines in untested environments, commonly referred to as CV1 (e.g. Burgueño et al. 2012; Mageto et al. 2020) in genomic prediction studies. For each test set, we defined prediction accuracy as the Pearson correlation between the predicted and observed phenotypic values of the lines.

## Abbreviations

GP: Genomic Prediction
GEBV: Genomic Estimated Breeding Values
CV: Cross Validation
GEI or G×E: Genotype-by-Environmental Interaction
BG-BLUP: Bayesian Genomic Best Linear Unbiased Prediction
CoV: Coefficient of Variation
SD: Standard Deviation
LD: Linkage Disequilibrium
LEN: Fibre Length
UNI: Fibre Uniformity
SFI: Short Fibre Index
STR: Fibre Strength
EL: Fibre Elongation
MIC: Micronaire
LY: Lint Yield
LP: Lint Percentage
EC: Environmental Covariate
DD: Day-Degree
ResNet: Residual Network
ENC: Encoder
FC: Fully Connected
BN: Batch Normalisation
ReLU: Rectified Linear Unit
VPD: Vapour Pressure Deficit

## Data availability

The phenotype, genotype, and environment data as well as the source code to implement the linkage equilibrium analysis, Bayesian regression and deep learning methods will be publicly available in the CSIRO Data Access Portal (https://data.csiro.au/) upon acceptance of the manuscript.

## Author contributions

ZL, WC and IW conceived the study questions and designed the research. WS, SL and WC developed and collected breeding lines, and conducted phenotype analysis. ZL and XL performed the genomic prediction analyses. ZL, WC, XL and QZ wrote the paper with contributions from all other authors.

## Acknowledgements

This study was financially supported by Cotton Breeding Australia, a joint venture between CSIRO and Cotton Seed Distributors Ltd. We are grateful for the technical expertise of Philippe Moncuquet, Dr Robert Davy, and Melanie Soliveres as well as the invaluable contribution to this work made by technical staff of the CSIRO cotton breeding group, who undertook the phenotyping and tissue sampling for genotyping. We also acknowledge Dr Pierce Rafter, Dr Qian Feng, Dr Angel Popa-Baez, and Dr Dung Ngoc Nguyen who contributed through general discussions and advice related to this project. Finally, we thank Dr Hien Nguyen for proofreading the manuscript.

## Declaration of interests

The authors declare no competing interests.

## Supplemental Information

**Table S1.**
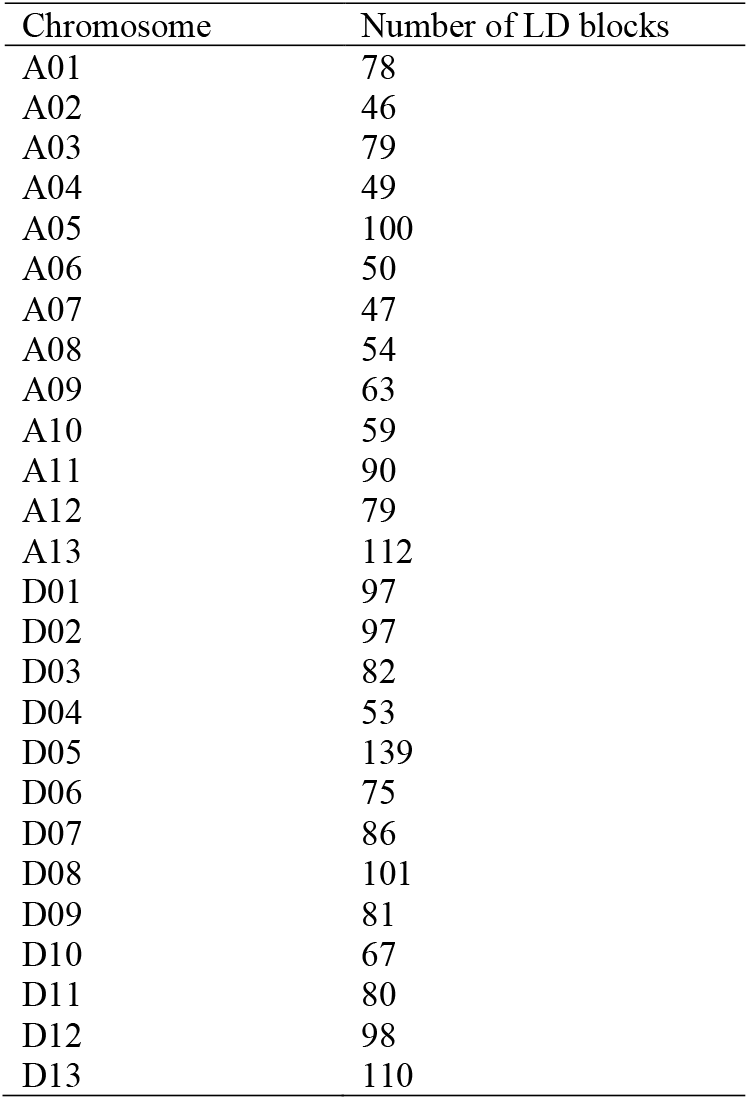
Distribution of LD blocks in each chromosome.

| Chromosome | Number of LD blocks |
| --- | --- |
| A01 | 78 |
| A02 | 46 |
| A03 | 79 |
| A04 | 49 |
| A05 | 100 |
| A06 | 50 |
| A07 | 47 |
| A08 | 54 |
| A09 | 63 |
| A10 | 59 |
| A11 | 90 |
| A12 | 79 |
| A13 | 112 |
| D01 | 97 |
| D02 | 97 |
| D03 | 82 |
| D04 | 53 |
| D05 | 139 |
| D06 | 75 |
| D07 | 86 |
| D08 | 101 |
| D09 | 81 |
| D10 | 67 |
| D11 | 80 |
| D12 | 98 |
| D13 | 110 |

**Table S2.**
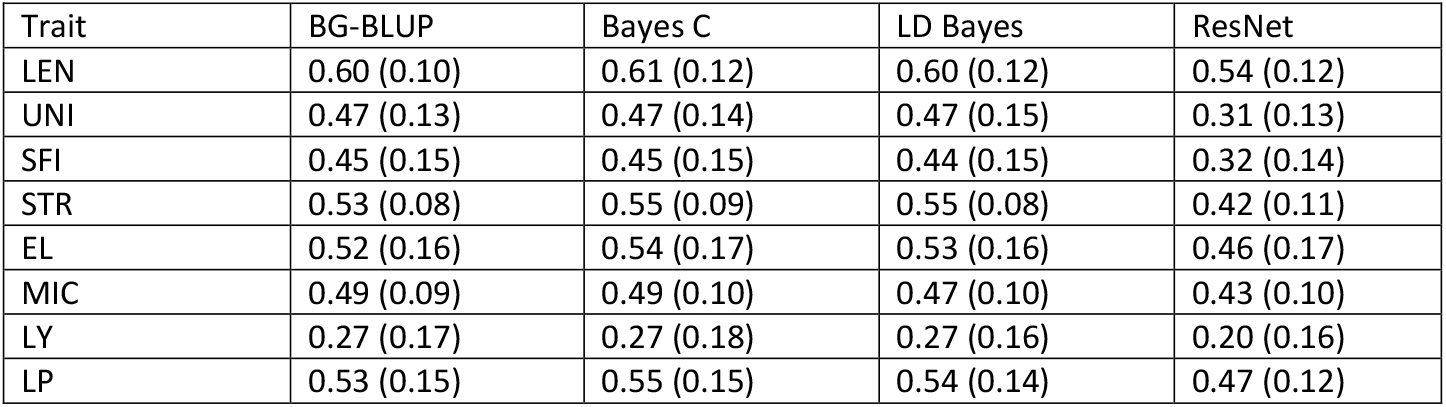
Mean prediction accuracies (standard error) over cross validation runs for eight traits including Fibre length, Uniformity, Short fibre index, Fibre Strength, Elongation, Micronaire, Lint yield and Lint percentage obtained using four genomic prediction methods (BG-BLUP, Bayes C, LD-Bayes, and ResNet).

**Table S3.** Mean prediction accuracies (standard error) across cross validation runs for eight traits including Fibre length, Uniformity, Short fibre index, Fibre Strength, Elongation, Micronaire, Lint yield and Lint percentage obtained using Bayes C, under the full G×E model, which includes additive genetic effects, environmental effects and their interactions. The G×E model is compared to two reduced models: (i) the G+E model, which only includes the additive effects of genetics and environments, and (ii) the G model which only includes the genetic effects.

| Trait | G model | G+E model | G×E model |
| --- | --- | --- | --- |
| LEN | 0.50 (0.12) | 0.57 (0.13) | 0.61 (0.12) |
| UNI | 0.35 (0.16) | 0.42 (0.15) | 0.47 (0.14) |
| SFI | 0.32 (0.15) | 0.40 (0.15) | 0.45 (0.15) |
| STR | 0.41 (0.15) | 0.50 (0.11) | 0.55 (0.09) |
| EL | 0.25 (0.20) | 0.49 (0.16) | 0.53 (0.16) |
| MIC | 0.36 (0.12) | 0.46 (0.09) | 0.47 (0.10) |
| LY | 0.17 (0.19) | 0.22 (0.16) | 0.27 (0.16) |
| LP | 0.39 (0.18) | 0.48 (0.14) | 0.55 (0.15) |

**Table S4.**
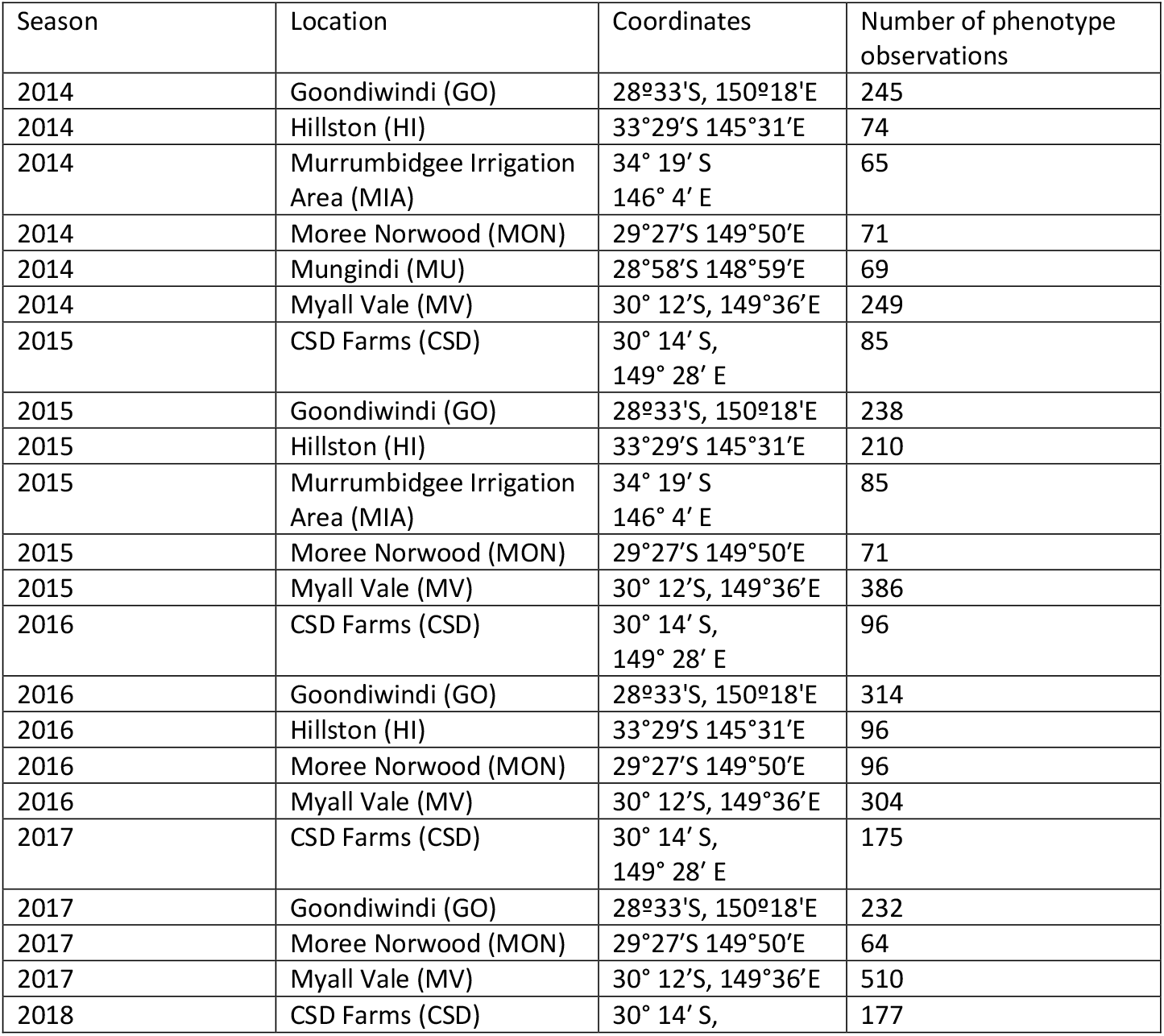
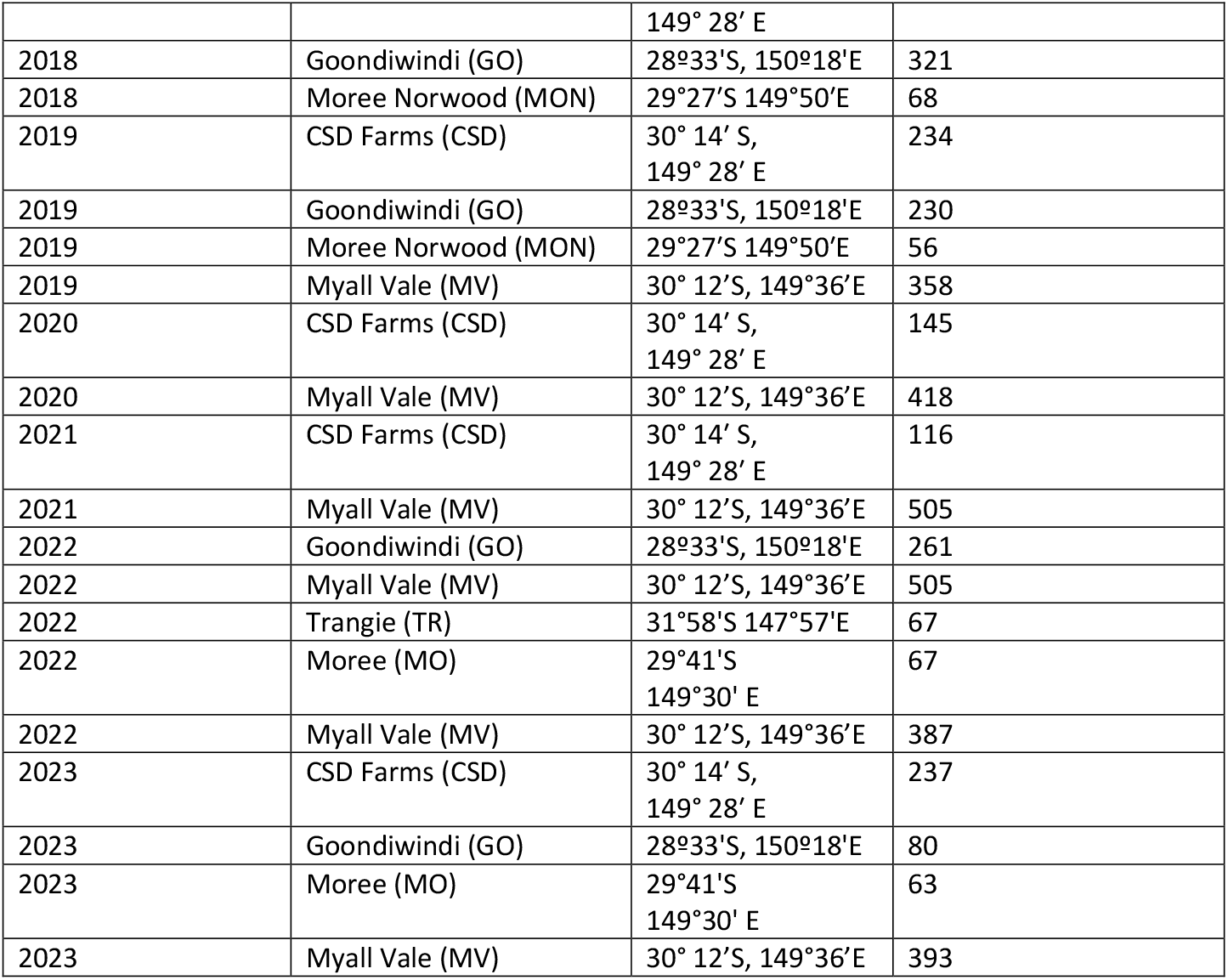
Locations and years where phenotypic data for genomic prediction analyses were collected.

**Table S5.** Cumulative day-degree thresholds (°C) associated with major cotton developmental stages, from first emergence through to crop maturity.

| Growth stages | Day degree targets (°C) |
| --- | --- |
| First emergence | 80 |
| First square | 505 |
| First flowering | 777 |
| Peak flowering | 1302 |
| Open boll | 1527 |
| Crop maturity | 2245 |

**Figure S1.**
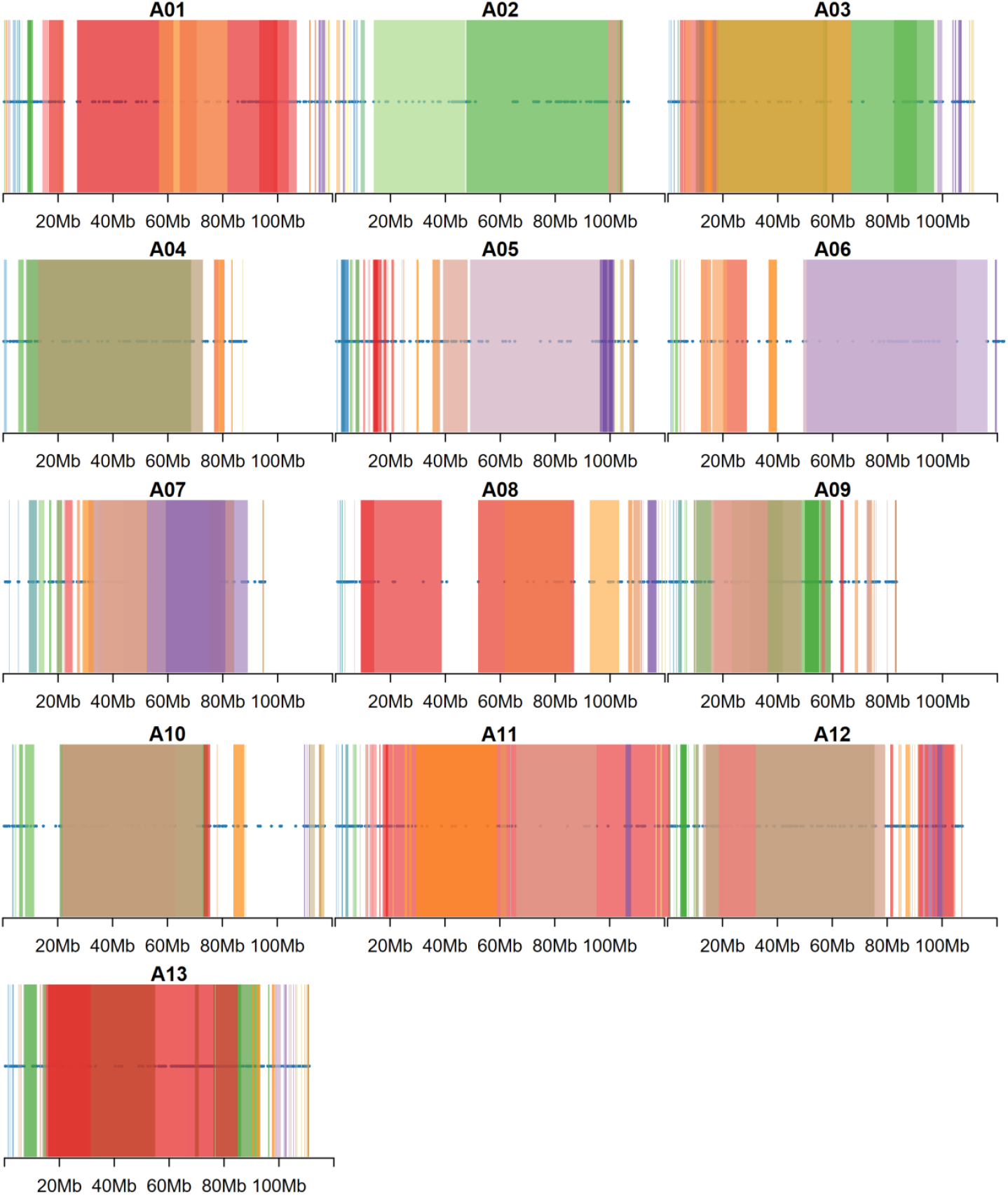
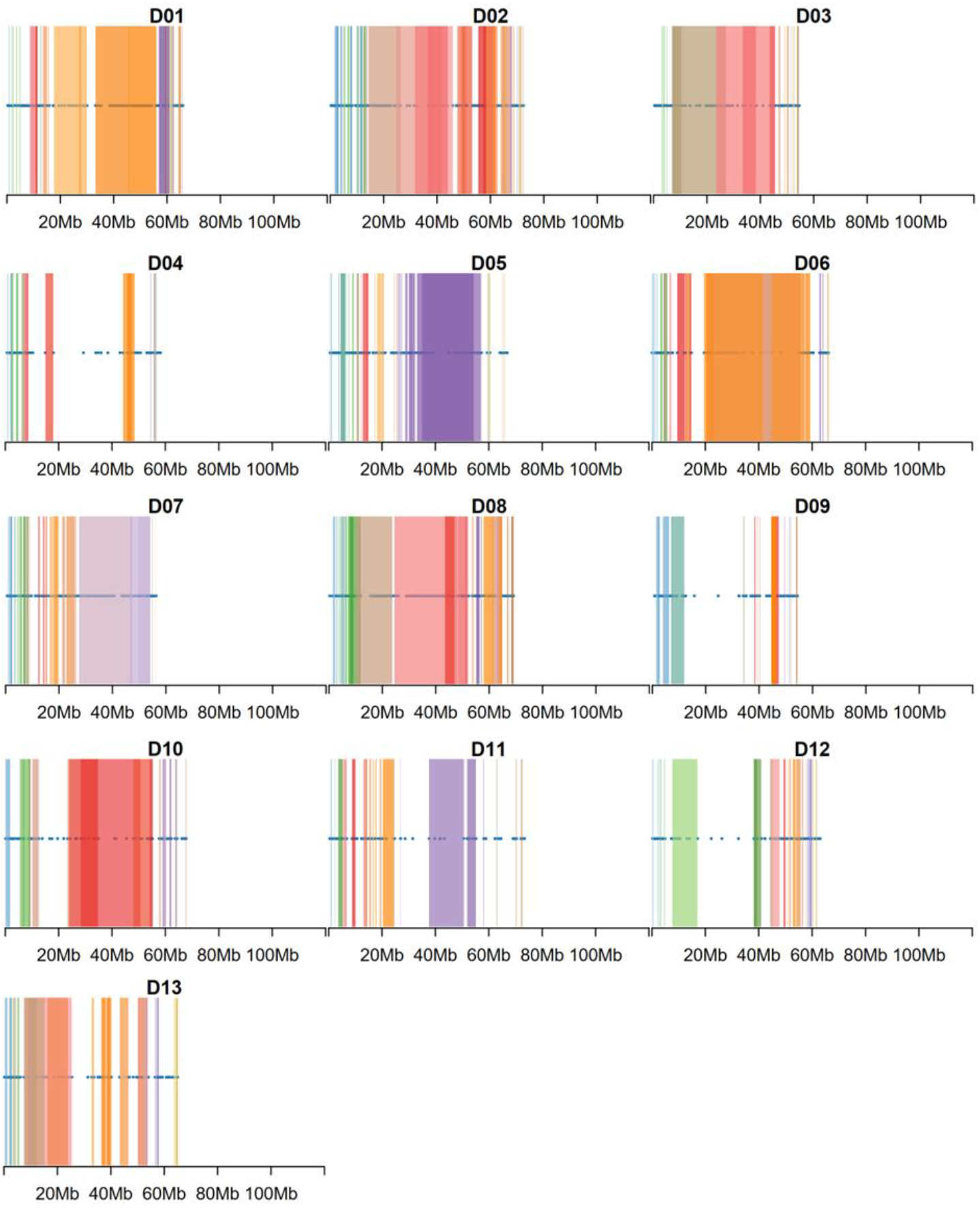
Linkage disequilibrium (LD) blocks plotted along each chromosome of the A-subgenome (A01–A13; top panel) and D-subgenome (D01–D13; bottom panel). Each coloured segment represents a contiguous LD block identified from SNP data, and horizontal genomic coordinates indicate physical positions (Mb).

**Figure S2.**
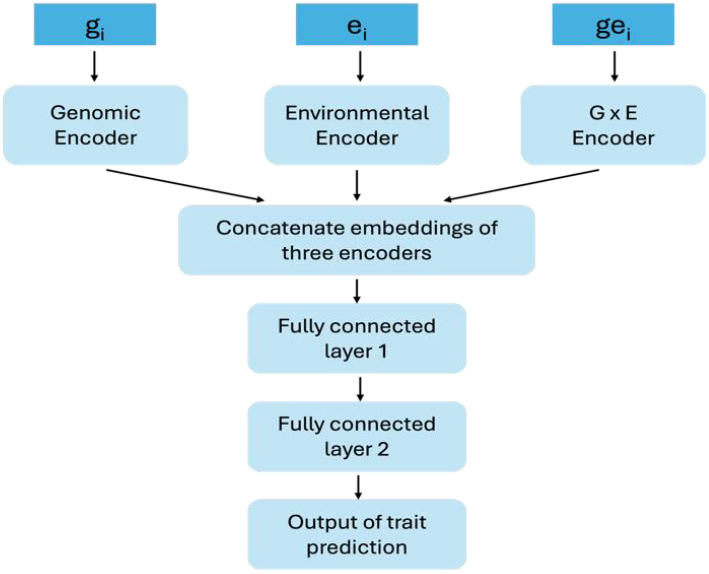
Framework of multimodal deep learning model Compared with single-modal or shallow baselines, this multimodal design explicitly models genomic, environmental and their interaction effects within a unified end-to-end framework.

## Notes

### Competing Interest Statement

The authors have declared no competing interest.

## References

Alemu, A., Åstrand, J., Montesinos-López, O.A., Sánchez, J.I., Fernández-Gónzalez, J., Tadesse, W., Vetukuri, R.R., Carlsson, A.S., Ceplitis, A., Crossa, J., Ortiz, R., and Chawade, A. (2024). Genomic selection in plant breeding: Key factors shaping two decades of progress. Mol Plant 17:552–578. 10.1016/j.molp.2024.03.007

Baghyalakshmi, K., Ariyapalayam, P.R., Govindaraj, S., Selvaraj, R., and Hiriyannaiah, P.A. (2024). Genetic improvement of fiber quality in tetraploid cotton: An overview of major QTLs and genes involved in and edited for the quality of cotton fibers. J Cotton Res 7:33. 10.1186/s42397-024-00196-9

Bange, M., Nunn, C., Mahan, J., Payton, P., Milroy, S., Finger, N., Caton, J., Dodge, W., and Quinn, J. (2022). Improving temperature-based predictions of the timing of flowering in cotton. J. Agron. 114:2728–2742. 10.1002/agj2.21086

Bernardo, R. (2008). Molecular markers and selection for complex traits in plants: learning from the last 20 years. Crop Sci. 48:1649–1664. 10.2135/cropsci2008.03.0131

Burgueño, J., de los Campos, G., Weigel, K., and Crossa, J. (2012) Genomic Prediction of Breeding Values when Modeling Genotype × Environment Interaction using Pedigree and Dense Molecular Markers. Crop Sci. 52:707–719. 10.2135/cropsci2011.06.0299

de los Campos, G., Sorensen, D., and Gianola, D. (2015). Genomic Heritability: What Is It? PLoS Genet. 11:e1005048. 10.1371/journal.pgen.1005048

Carvalho, H.F., Rio, S., Garcia-Abadillo, J., and Sánchez, J.I. (2024). Revisiting superiority and stability metrics of cultivar performances using genomic data: derivations of new estimators. Plant Methods 20:85. 10.1186/s13007-024-01231-1

Chen, Y., Gao, Y., Chen, P.Y., Zhou, J., Zhang, C.Y., Song, Z.Q., Huo, X.H., Du, Z.H., Gong, J.W., Zhao, C.J., Wang, S.L., Zhang, J.X., Wang, F.R., and Zhang, J. (2022). Genome-wide association study reveals novel quantitative trait loci and candidate genes of lint percentage in upland cotton based on the CottonSNP80K array. Theor. Appl. Genet. 135: 2279–2295. 10.1007/s00122-022-04111-1

Cobb, J.N., Biswas, P.S., and Platten, J.D. (2019). Back to the future: revisiting MAS as a tool for modern plant breeding. Theor. Appl. Genet. 132:647–667. 10.1007/s00122-018-3266-4

Conaty, W.C., Mahan, J.R., Neilsen, J.E., and Constable, G.A. (2014). Vapour pressure deficit aids the interpretation of cotton canopy temperature response to water deficit. Funct. Plant Biol. 41:535–546. 10.1071/FP13223

Constable, G.A., and Shaw, A.J. (1988). Temperature requirements for cotton (1^st^ ed., Agfact P5.3.5). NSW Agric and Fisheries, Orange, NSW, Australia.

Cooper, M., van Eeuwijk, F.A., Hammer, G.L., Podlich, D.W., and Messina, C. (2009). Modeling QTL for complex traits: Detection and context for plant breeding. Curr. Opin. Plant Biol. 12:231–240. 10.1016/j.pbi.2009.01.006

Costa-Neto, G., Gali, G., R., Carvalho, H.F., Crossa, J., and Fritsche-Neto, R. (2021). EnvRtype: a software to interplay enviromics and quantitative genomics in agriculture. G3 Genes Genomes Genet. 11:jkab040. 10.1093/g3journal/jkab040

Cotton Research and Development Corporation (2025). Australian Cotton Production Manual. https://crdc.com.au/sites/default/files/pdf/2025%20Australian%20Cotton%20Production%20Manual_interactive_sml.pdf.

Crossa, J., Fritsche-Neto, R., Montesinos-Lopez, O.A., Costa-Neto, G., Dreisigacker, S., Montesinos-Lopez, A., and Bentley, A.R. (2021). The modern plant breeding triangle: Optimizing the use of genomics, phenomics, and enviromics data. Front Plant Sci 12:651480. 10.3389/fpls.2021.651480

Crossa, J., Montesinos-López, O.A., Pérez-Rodríguez, P., Costa-Neto, G., Fritsche-Neto, R., Ortiz, R., Martini, J.W.R., Lillemo, M., Montesinos-López, A., Jarquin, D., Breseghello, F., Cuevas, J., and Rincent, R. (2022) Genome and environment based prediction models and methods of complex traits incorporating genotype × environment interaction. In: Montesinos-López OA, Montesinos-López A (eds) Genomic Prediction of Complex Traits: Methods and Protocols. Methods in Molecular Biology, vol 2467. Springer, New York, pp 245–283. 10.1007/978-1-0716-2205-6_9

Crossa, J., Pérez-Rodríguez, P., Cuevas, J., Montesinos-López, O., Jarquín, D., de los Campos, G., Burgueño, J., González-Camacho, J.M., Pérez-Elizalde, S., Beyene, Y., Dreisigacker, S., Singh, R., Zhang, X., Gowda, M., Roorkiwal, M., Rutkoski, J., and Varshney, R.K. (2017). Genomic selection in plant breeding: methods, models, and perspectives. Trends Plant Sci 22:961–975. 10.1016/j.tplants.2017.08.011

Crossa, J., Martini, J.W.R., Vitale, P., Pérez-Rodríguez, P., Costa-Neto, G., Fritsche-Neto, R., Runcie, D., Cuevas, J., Toledo, F., Li, H., De Vita, P., Gerard, G., Dreisigacker, S., Crespo-Herrera, L., Saint Pierre, C., Bentley, A., Lillemo, M., Ortiz, R., Montesinos-López, O.A., and Montesinos-López, A. (2025). Expanding genomic prediction in plant breeding: harnessing big data, machine learning, and advanced software. Trends Plant Sci. 30:756–774. 10.1016/j.tplants.2024.12.009

Doerge, R.W. (2002). Mapping and analysis of quantitative trait loci in experimental populations. Nat. Rev. Genet. 3:43–52. 10.1038/nrg703

Endelman, J.B. (2011). Ridge regression and other kernels for genomic selection with R package rrBLUP. Plant Genome 4:250–255. 10.3835/plantgenome2011.08.0024

Fang, D.D., Zeng, L., Thyssen, G.N., Delhom, C.D., Bechere, E., Jones, D.C., and Li, P. (2021). Stability and transferability assessment of the cotton fiber strength QTL qFS-c7-1 on chromosome A07. The Crop Journal 9:380–386. 10.1016/j.cj.2020.06.016

Fraley C., Raftery A.E., Murphy T.B., and Scrucca L. (2012). mclust Version 4 for R: Normal Mixture Modeling for Model-Based Clustering, Classification, and Density Estimation. Technical Report No. 597, Department of Statistics, University of Washington. http://www.dpye.iimas.unam.mx/lety/archivos/cursoinegi/INEGI%20CURSO%203-7%20AGOSTO2015/cluster/notasmclust.pdf

Gapare, W., Liu, S., Conaty, W., Zhu, Q-H., Gillepie, V., Llewellyn, D., Stiller, W., and Wilson, I. (2018). Historical datasets support genomic selection models for the prediction of cotton fiber quality phenotypes across multiple environments. Genes Genomes Genet. 8:1721–1732.

Goodfellow, I., Bengio, Y., and Courville, A. (2016). Deep Learning. MIT Press, Cambridge, MA.

Habier, D., Fernando, R.L., Kizilkaya, K., and Garrick D.J. (2011). Extension of the Bayesian alphabet for genomic selection. BMC Bioinformatics 12:186. 10.1186/1471-2105-12-186

Hajheidari, M., Sunyaev, S., and de Meaux, J. (2024) Are complex traits underpinned by polygenic molecular traits? A reflection on the complexity of gene expression. Plant Cell Physiol. 66:444–460. 10.1093/pcp/pcae140

Halpin-McCormick, A., Campbell, Q., Negrão, S., Morrell, P.L., Hübner, S., Neyhart, J.L., and Kantar, M.B. (2025). Environmental genomic selection to leverage polygenic local adaptation in barley landraces. Commun. Biol. 8:618. 10.1038/s42003-025-08045-4

He, K., Zhang, X., Ren, S., and Sun, J. (2016). Deep residual learning for image recognition. In Proceedings of the IEEE conference on computer vision and pattern recognition, pp. 770-778.

Heilmann, P.G., Grosch, E., Frisch, M., Herrmann, M., Beuch, S., Kurra, V., Mascher, M., Avni, R., Oldach, K., Röhrs, I., Hanemann, A., Mehta, R.R., Reinbrecht, C., Serfling, A., Stahl, A., Stucke, M., Abbadi, A., Kox, T., and Zenke-Philippi, C. (2025). Haplotype-based autoencoders can reduce the dataset dimension and estimate haplotype block effects in different crop species. BMC Bioinformatics 26:289. 10.1186/s12859-025-06323-w

Ishwaran, H., and Rao, J.S. (2005). Spike and Slab variable selection: frequentist and Bayesian strategies. Ann. Stat. 33:730–773. 10.1214/009053604000001147

Islam, M.S., Fang, D.D., Jenkins, J.N., Guo, J., McCarty, J.C., and Jones, D.C. (2020). Evaluation of genomic selection methods for predicting fiber quality traits in Upland cotton. Mol. Genet. Genom. 295:67–79. 10.1007/s00438-019-01599-z

Jabran, K., Ul-Allah, S., Chauhan, B.S., and Bakhsh, A. (2019). An introduction to global production trends and uses, history and evolution, and genetic and biotechnological improvements in cotton. In: Jabran K, Chauhan BS Eds. Cotton Production. Wiley, Hoboken, UK, p 1–22.

Jannink, J.L., Lorenz, A.J., and Iwata, H. (2010). Genomic selection in plant breeding: from theory to practice. Brief Funct Genomics 9:166–177. 10.1093/bfgp/elq001

Jarquín, D., Crossa, J., Lacaze, X., Du Cheyron, P., Daucourt, J., Lorgeou, J., Piraux, F., Guerreiro, L., Pérez, P., Calus, M., Burgueño, J., and de los Campos, G. (2014). A reaction norm model for genomic selection using high dimensional genomic and environmental data. Theor. Appl. Genet. 127:595–607. 10.1007/s00122-013-2243-1

Jeffrey, S.J., Carter, J.O., Moodie, K.B., and Beswick, A.R. (2001). Using spatial interpolation to construct comprehensive archive of Australian climate data. Environ. Model Softw. 16:309–330. 10.1016/S1364-8152(01)00008-1

Kemppainen, P., Knight, C.G., Sarma, D.K., Hlaing, T., Prakash, A., MaungMaung, Y.N., Somboon, P., Mahanta, J., and Walton, C. (2015) Linkage disequilibrium network analysis (LDna) gives a global view of chromosomal inversions, local adaptation and geographic structure. Mol. Ecol. Res. 15:1031–104. 10.1111/1755-0998.12369

Kingma, D.P., and Ba, J. (2014). Adam: A method for stochastic optimization. arXiv preprint 1412.6980.

Li, Z., Kemppainen, P., Rastas. P., and Merilä, J. (2018). Linkage disequilibrium clustering-based approach for association mapping with tightly linked genome wide data. Mol. Ecol. Res. 18:809–824. 10.1111/1755-0998.12893

Li, Z., Liu, S., Conaty, W., Zhu, Q-H., Moncuquet, P., Stiller, W., and Wilson, I. (2022). Genomic prediction of cotton fibre quality and yield traits using Bayesian regression methods. Heredity 129:103–112. 10.1038/s41437-022-00537-x.

Li, Z., Zhu, Q-H., Moncuquet, P., Wilson, I., Llewellyn, D., Stiller, W., and Liu, S. (2024). Quantitative genomics-enabled selection for simultaneous improvement of lint yield and seed traits in cotton (Gossypium hirsutum L.). Theor. Appl. Genet. 137:142. 10.1007/s00122-024-04645-6.

Liu, S.M., Llewellyn, D.J., Stiller, W.N., Jacobs, J., Lacape, J.M., and Constable, G.A. (2011). Heritability and predicted selection response of yield components and fibre properties in an inter-specific derived RIL population of cotton. Euphytica 178:309–320.

Liu, S.M., Constable, G.A., Cullis, B.R., Stiller, W.N., and Reid, P.E. (2015). Benefit of spatial analysis for furrow irrigated cotton breeding trials. Euphytica 201:253–264. 10.1007/s10681-014-1205-2

Liu, S.M., and Constable, G.A. (2017). Effect of self-generation for initial selection on breeding better cotton. Euphytica 213:17. 10.1007/s10681-017-2052-8

Lourenço, V.M., Ogutu, J.O., Rodrigues, R.A.P., Posekany, A., and Piepho, H.P. (2024). Genomic prediction using machine learning: a comparison of the performance of regularized regression, ensemble, instance-based and deep learning methods on synthetic and empirical data. BMC Genomics 25:152. 10.1186/s12864-023-09933-x

Mageto, E.K., Crossa, J., Pérez-Rodríguez, P., Dhliwayo, T., Palacios-Rojas, N., Lee, M., Guo, R., San Vicente, F., Zhang, X., and Hindu, V. (2020). Genomic Prediction with Genotype by Environment Interaction Analysis for Kernel Zinc Concentration in Tropical Maize Germplasm. G3 Genes Genomes Genet. 10:2629–2639. 10.1534/g3.120.401172

Manthena, V., Jarquín, D., Varshney, R.K., Roorkiwal, M., Dixit, G.P., Bharadwaj, C., and Howard, R. (2022). Evaluating dimensionality reduction for genomic prediction. Front. Genet. 13:958780. 10.3389/fgene.2022.958780

Meher, P.K., Kumar, A., and Pradhan, S.K. (2022). Genomic Selection Using Bayesian Methods: Models, Software, and Application. In: Genomics of Cereal Crops. Springer Protocols Handbooks, pp 259–269. Springer, New York. 10.1007/978-1-0716-2533-0_13

Meuwissen, T.H., Hayes, B.J., and Goddard, M.E. (2001). Prediction of total genetic value using genome-wide dense marker maps. Genetics 157:1819–1829. 10.1093/genetics/157.4.1819

Meyer, K. (2009). Factor-analytic models for genotype × environment type problems and structured covariance matrices. Genet Sel Evol 41:21. 10.1186/1297-9686-41-21

Montesinos-López, O.A., Montesinos-López, A., Pérez-Rodríguez, P., Barrón-López, J.A., Martini, J.W.R., Fajardo-Flores, S.B., Gaytan-Lugo, L.S., Santana-Mancilla, P.C., and Crossa, J. (2021). A review of deep learning applications for genomic selection. BMC Genomics 22:19. 10.1186/s12864-020-07319-x

Montesinos-López, A., Rivera, C., Pinto, F., Piñera, F., Gonzalez, D., Reynolds, M., Pérez-Rodríguez, P., Li, H., Montesinos-López, O.A., and Crossa, J. (2023) Multimodal deep learning methods enhance genomic prediction of wheat breeding. G3 Genes Genomes Genet 13:jkad045. 10.1093/g3journal/jkad045

Montesinos-López, O.A., Chavira-Flores, M., Kismiantini Crespo-Herrera, L., Saint Pierre, C, Li, H., Fritsche-Neto, R., Al-Nowibet, K., Montesinos-López, A., and Crossa, J. (2024) A review of multimodal deep learning methods for genomic-enabled prediction in plant breeding. Genetics 228:iyae161. 10.1093/genetics/iyae200

Napier, J.D., Heckman, R.W., and Juenger, T.E. (2023). Gene-by-environment interactions in plants: Molecular mechanisms, environmental drivers, and adaptive plasticity. Plant Cell 35: 109–124. 10.1093/plcell/koac322

Ngiam, J., Khosla, A., Kim, M., Nam, J., Lee, H., and Ng, A,Y. (2011) Multimodal deep learning. In: Proceedings of the 28th International Conference on Machine Learning (ICML); Bellevue, WA, USA. p. 689–696.

Niu, H., Ge, Q., Shang, H., Yuan, Y. (2022). Inheritance, QTLs, and candidate genes of lint percentage in upland cotton. Front. Genet. 13:855574. 10.3389/fgene.2022.855574

O’Hara, R.B., and Sillanpää, M.J. (2009). A review of Bayesian variable selection methods: what, how and which. Bayesian Anal. 4:85–117. 10.1214/09-BA403

Paterson, A.H., Wendel, J.F., Gundlach, H, Guo, H., Jenkins, J., Jin, D., Llewellyn, D., Showmaker, K.C., Shu, S., Udall, J. et al. (2012) Repeated polyploidization of Gossypium genomes and the evolution of spinnable cotton fibres. Nature 492:423–427. 10.1038/nature11798

Pérez, P., and de los Campos, G. (2014) Genome-wide regression and prediction with the bglr statistical package. Genetics 198:483–495. 10.1534/genetics.114.164442

Scarpin, G.J., Bhattarai, A., Hand, L.C., Snider, J.L., Roberts, P.M., and Bastos, L.M. (2025). Cotton lint yield and quality variability in Georgia, USA: understanding genotypic and environmental interactions. Field Crops Res. 325:109822. 10.1016/j.fcr.2025.109822

Shang, L., Wang, Y., Wang, X., Liu, F., Abduweli, A., Cai, S., Li, Y., Ma, L., Wang, K., and Hua, J. (2016). Genetic analysis and stable QTL detection on fiber quality traits using two recombinant inbred line populations and their backcross progeny in upland cotton. G3 Genes Genomes Genet 6:2717–2724. 10.1534/g3.116.032359

Sharif, I., Aleem, S., Junaid, J.A., Aleem, M., Jamshaid, K., Saleem, H., Rizwan, M., Chohan, S.M., Sohail, S., Akram, S., Zeeshan, M., and Sarwar, G. (2025). Evaluation of genotype × environment interaction and yield stability of Cotton (Gossypium hirsutum L.) genotypes under heat stress conditions. J. Crop Health 77:16. 10.1007/s10343-024-01079-4

Shen, C., Li, X., Zhang, R., and Lin, Z. (2017) Genome-wide recombination rate variation in a recombination map of cotton. PLoS One 12: e0188682. 10.1371/journal.pone.0188682

Snider, J.L., Collins, G.D., Whitaker, J., and Davis, J.W. (2013). Quantifying genotypic and environmental contributions to yield and fiber quality in Georgia: data from seven commercial cultivars and 33 yield environments. J. Cotton Sci. 17:285–292. https://www.cotton.org/journal/2013-17/4/upload/JCS17-285.pdf

Song, H., Dong, T., Wang, W., Yan, X., Geng, C., Bai, S., and Hu, H. (2025). A deep autoencoder compression-based genomic prediction method for whole-genome sequencing data. Biology 14:89. 10.3390/biology14111622

Sreedasyam, A., Lovell, J.T., Mamidi, S., Khanal, S., Jenkins, J.W., Plott, C., Bryan, K.B., Li, Z., Shu, S., Carlson, J. et al. (2024). Genome resources for three modern cotton lines guide future breeding efforts. Nat. Plants 10:1039–1051. 10.1038/s41477-024-01713-z

Stekhoven, D.J., and Bühlmann, P. (2012). MissForest—non-parametric missing value imputation for mixed-type data. Bioinformatics 28:112–118. 10.1093/bioinformatics/btr597

Tolhurst, D.J., Gaynor, R.C., Gardunia, B., Hickey, J.M., and Gorjanc, G. (2022). Genomic selection using random regressions on known and latent environmental covariates. Theor. Appl. Genet. 135:3393–3415. 10.1007/s00122-022-04186-w

Wang, M., Tu, L., Lin, M., Lin, Z., Wang, P., Yang, Q., Ye, Z., Shen, C., Li, J., Zhang, L. et al. (2017). Asymmetric subgenome selection and cis-regulatory divergence during cotton domestication. Nat. Genet. 49:579–587. 10.1038/ng.3807

Wang, K., Abid, M.A., Rasheed, A., Crossa, J., Hearne, S., and Li, H. (2023). DNNGP, a deep neural network-based method for genomic prediction using multi-omics data in plants. Mol. Plant 16:279–293. 10.1016/j.molp.2022.11.004

VanRaden, P.M. (2008). Efficient methods to compute genomic predictions. J. Dairy Sci. 91:4414–4423. 10.3168/jds.2007-0980

Vaswani, A., Shazeer, N., Parmar, N., Uszkoreit, J., Jones, L., Gomez, A.N., Kaiser, Ł., and Polosukhin, I. (2017). Attention Is All You Need. Advances in Neural Information Processing Systems (NIPS 2017). https://arxiv.org/pdf/1706.03762

Virk, G., Snider, J.L., and Bourland, F.M. (2023). Genotypic and environmental contributions to lint yield, yield components, and fiber quality in upland cotton from Arkansas variety trials over a 19-year period. Crop Sci. 63:1284–1299. 10.1002/csc2.20948

Voss-Fels, K.P., Cooper, M., and Hayes, B.J. (2019) Accelerating crop genetic gains with genomic selection. Theor. Appl. Genet. 132:669–686. 10.1007/s00122-018-3120-8

Washburn, J.D., Cimen, E., Ramstein, G., Reeves, T., O’Briant, P., McLean, G., Cooper, M., Hammer, G., and Buckler, E.S. (2021). Predicting phenotypes from genetic, environment, management, and historical data using CNNs. Theor. Appl. Genet. 134:3997–4011. 10.1007/s00122-021-03943-7

Xiong, W., Reynolds, M., and Xu, Y. (2022) Climate change challenges plant breeding. Curr. Opin. Plant Biol. 70: 102308. 10.1016/j.pbi.2022.102308

Yao, Z., Yao, M., Wang, C., Li, K., Guo, J., Xiao, Y., Yan, J., and Liu, J. (2025). GEFormer: A genotype–environment interaction–based genomic prediction method that integrates the gating multilayer perceptron and linear attention mechanisms. Mol. Plant 18, 527–549. 10.1016/j.molp.2025.01.020

Ye, H., Zhang, Z., Ren, D., Cai, X., Zhu, Q., Ding, X., Zhang, H., Zhang, Z., and Li, J. (2022). Genomic Prediction Using LD-Based Haplotypes in Combined Pig Populations. Front. Genet. 13:843300. 10.3389/fgene.2022.843300

You, J., Ellis, J.L., Adams, S., Sahar, M., Jacobs, M., and Tulpan, D. (2023) Comparison of imputation methods for missing production data of dairy cattle. Animal 17:100921. 10.1016/j.animal.2023.100921

Zhang, Q., Privé, F., Vilhjálmsson, B., and Speed, D. (2021). Improved genetic prediction of complex traits from individual-level data or summary statistics. Nat. Commun. 12:4192. 10.1038/s41467-021-24485-y

Zhang, D., Yang, F., Li, J., Liu, Z., Han, Y., Zhang, Q., Pan, S., Zhao, X., and Wang, K. (2025). Progress and perspectives on genomic selection models for crop breeding. Technol. Agron. 5:e006. 10.48130/tia-0025-0002

